# Gut microbiota-derived indole-3-propionic acid preserves dorsal hippocampal catecholamines to prevent post-stroke cognitive impairment

**DOI:** 10.64898/2026.04.27.721133

**Authors:** Yisi Liu, Huaying Zhang, Fangbo Xia, Xuxuan Gao, Zhuang Li, Xin Zhao, Fan Wu, Mengxi Li, Kaiyu Xu, Muxuan Chen, Ranyue Ren, Weike Hu, Jia Yin, Hongwei Zhou, Dongxin Zhang

## Abstract

**Background:** Gut dysbiosis has been increasingly implicated in post-stroke cognitive impairment (PSCI), yet the causal contribution and therapeutic potential of gut microbiota-derived metabolites remain unclear. This study aimed to identify key microbiota-derived metabolites involved in PSCI and to elucidate their underlying mechanisms.

**Results:** We found that both PSCI patients and middle cerebral artery occlusion (MCAO) mice exhibited distinct gut microbial alterations, characterized by a marked reduction in tryptophan-metabolizing bacteria and indole-3-propionic acid (IPA), a gut microbiota-derived tryptophan metabolite. Exogenous IPA administration alleviated PSCI-like phenotypes in MCAO mice. Mechanistically, IPA preserved tyrosine hydroxylase-positive (Th⁺) fibers and catecholamine levels in the dorsal hippocampus. Further analyses showed that IPA binds to the adaptor protein Ywhab, promotes ERK activation, and enhances neuronal survival, thereby counteracting neuronal apoptosis-associated inflammation and subsequent Th⁺ fiber degeneration.

**Conclusion:** These findings identify IPA as a gut microbiota-derived neuromodulator that mitigates PSCI by preserving dorsal hippocampal catecholaminergic transmission. IPA may therefore serve as a promising predictive biomarker and therapeutic candidate for PSCI.

## Introduction

Cognitive impairment, which affects approximately 38% of stroke survivors, represents a significant health challenge, and adversely impacts long-term recovery after stroke^1–3^. The pathogenesis of PSCI (post-stroke cognitive impairment) is influenced by infarct characteristics, cardiovascular risk factors, and brain resilience^4^. Current pharmacological interventions for PSCI are largely based on those used for neurodegenerative diseases or vascular cognitive impairment, with acetylcholinesterase inhibitors being the most commonly used agents. Yet, the effectiveness of these interventions remains limited, underscoring the need for exploring its pathogenesis and developing targeted therapeutic strategies^4^.

Accumulating evidence highlights the role of gut microbiota-derived metabolites in modulating central nervous system function and pathology, such as promoting nerve regeneration^6^ and alleviating neuroinflammation^7^. Although gut dysbiosis has been implicated in PSCI^8–11^, the specific metabolites derived from gut microbiota, their temporal changes after stroke, and the mechanisms by which they exert effects have yet to be fully delineated.

Gut bacterial metabolism of dietary tryptophan produces a diverse array of indole derivatives that serve as key molecular mediators of gut-brain communication^12,13^. These indole metabolites activate receptors such as aryl hydrocarbon receptor and pregnane X receptor, thereby shaping neuroinflammatory pathways and oxidative stress responses^7,14^. Through these mechanisms, tryptophan-derived indole metabolites exert potent neuroprotective and anti-inflammatory actions^7,15^. However, current research focused largely on their effects during the acute phase of stroke, leaving their long-term roles in neuronal remodeling and cognitive recovery insufficiently understood^16^. As a result, the mechanisms by which they influence delayed neurodegeneration and sustained cognitive impairment after stroke remain unclear.

Among these indole metabolites, indole-3-propionic acid (IPA) stands out as a potent and abundant compound with antioxidative and metabolic benefits^16^. Previous studies have demonstrated that IPA exerts potent protective effects in conditions such as diabetes^19^, hyperlipidemia^20^, and hypertension^21^, that are major risk factors for stroke. In addition, IPA has been shown to reduce infarct volume in experimental models of ischemic stroke^21^. Nevertheless, whether IPA contributes to post-stroke cognitive recovery and the specific mechanisms have not been elucidated.

This study shows that gut tryptophan-metabolizing bacteria and IPA are significantly reduced in both MCAO (middle cerebral artery occlusion) mice and stroke patients, particularly in those who develop PSCI. Thus, we investigated whether IPA could ameliorate PSCI, and uncovered specific molecular mechanism underlying its role in neuroprotection and cognitive recovery following stroke.

## Results

### The reduction of gut microbiota-derived IPA in both PSCI patients and MCAO mice

We enrolled a cohort of 84 patients with stroke, and stratified them into those with and without post-stroke cognitive impairment (PSCI and nPSCI) at the 3-month follow-up, based on the Montreal Cognitive Assessment (MoCA) using a cutoff score of 26^23,24^. Patients who developed PSCI exhibited significantly altered gut microbiota composition during the acute phase, with notable differences in both α- and β-diversity compared to nPSCI patients (Fig. 1a and Fig. S1a-d). To further identify gut bacterial features associated with cognitive impairment, LEfSe analysis was performed on gut microbiota during the acute phase. The analysis revealed a pronounced reduction in several beneficial taxa, including *Bifidobacteriales*, *Lactobacillales* and certain *Clostridiales*, in patients who later developed PSCI (Fig. 1b). These gut bacteria are typically involved in tryptophan metabolism, producing a range of neuroactive indole derivatives^25–28^. At the 90-day follow-up, persistent gut dysbiosis occurred in the PSCI group, characterized by reduced abundances of *Bifidobacteriales* and *Lactobacillales* compared to nPSCI group (Fig. 1c-d and Fig. S1e-h). We next profiled the serum metabolome to uncover the differential gut-derived tryptophan metabolites (Fig. 1e). Among the candidate metabolites identified, IPA, a neuroprotective tryptophan metabolite produced by the gut microbiota^29,30^, emerged as one of the significantly depleted metabolites in PSCI patients during the acute phase (Fig. 1f-g). Notably, this IPA deficiency persisted throughout the 90-day follow-up period (Fig. 1h), suggesting a sustained disruption in gut bacterial biosynthesis of IPA in individuals vulnerable to cognitive impairment. These findings suggest that impaired production of IPA may link gut microbiota dysbiosis and PSCI, highlighting IPA as a potential therapeutic target for PSCI.

**Fig. 1.**
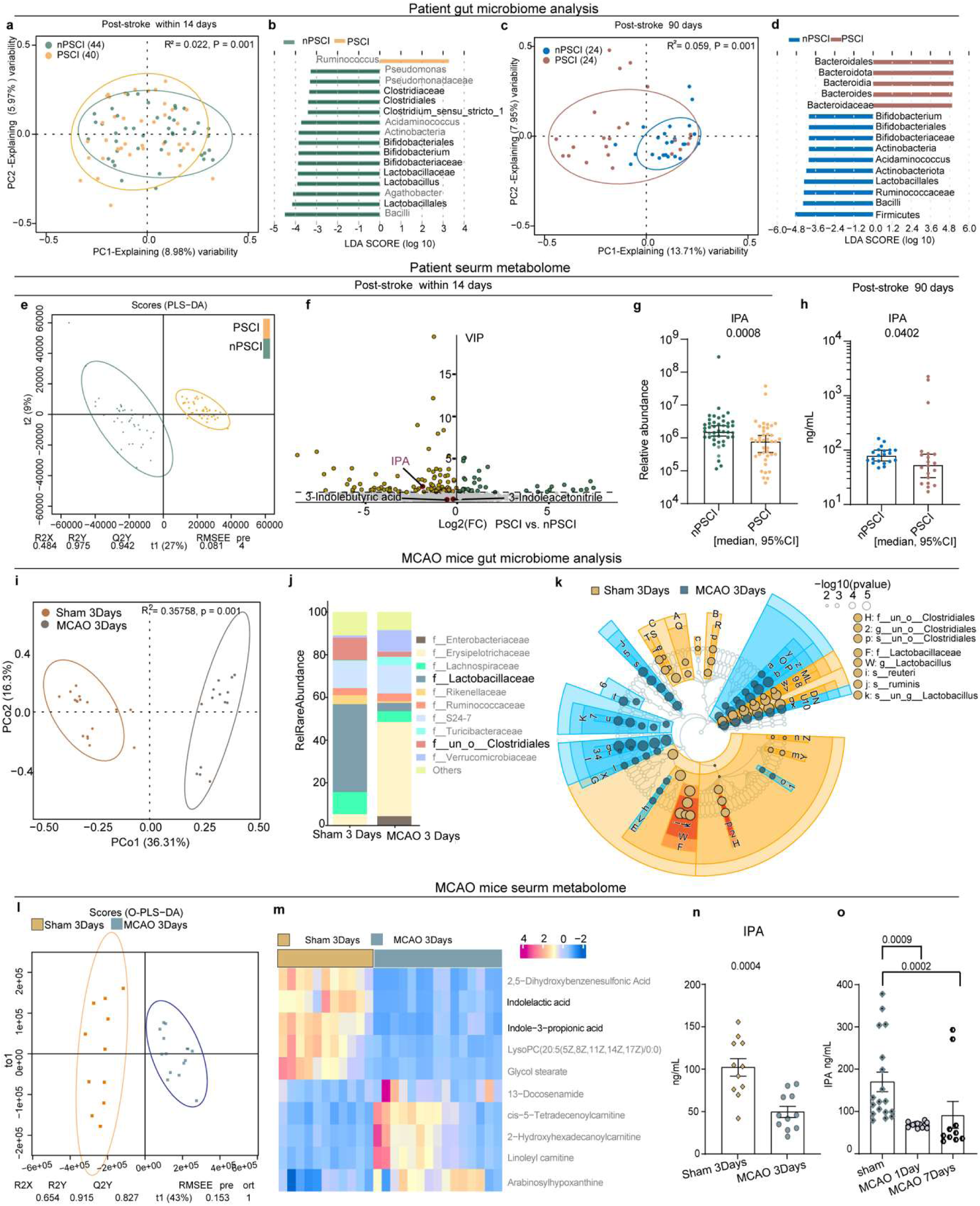
The reduction of gut microbiota-derived IPA in both PSCI patients and MCAO mice. (a-d) The 16S rRNA sequencing analysis of gut microbiota in stroke patients. (a, c) PCoA plots of β-diversity based on Bray-Curtis distances, showing distinct microbial community structures between nPSCI and PSCI groups during the acute phase (a) and at follow-up (c). (b, d) LEfSe analysis identifying differentially abundant taxa between the two groups during the acute phase (b) and at follow-up (d). (e) Metabolomics analysis of serum from patients with or without PSCI (n = 44, 40). (f) Differentially expressed metabolites between the two groups during the acute phase, identified based on VIP > 1. (g) Relative abundance of indole-3-propionic acid (IPA) in PSCI and nPSCI groups during the acute phase (n = 44, 40). (h) Serum concentrations of IPA in PSCI and nPSCI groups at follow-up (n = 20 per group). (i) Principal Coordinates Analysis (PCoA) of gut bacterial composition between Sham and MCAO groups at 3 days post-stroke (n = 14, 17). (j) Gut bacterial composition at the family level in Sham and MCAO groups at 3 days post-stroke (n = 14, 17). (k) LEfSe analysis identifying differentially abundant taxa between Sham and MCAO groups. (l) Serum metabolic profiles of Sham and MCAO groups at 3 days post-stroke (n = 11, 15). (m) The top 10 metabolites upregulated and downregulated in serum of mice that underwent MCAO or Sham. (n) The serum levels of IPA in Sham and MCAO groups at 3 days post-stroke (n = 11 per group). (o) The levels of IPA in the serum of mice following 1 and 7 days of MCAO or Sham (n = 18, 12, 10). M-W test (g-h), Unpaired t test (n). K-W test with Dunn’s test (o). The results are expressed as [median, 95%CI] (g-h). The results are expressed as means ± s.e.m. (n-o).

Moreover, we employed MCAO (middle cerebral artery occlusion) mice model to assess the changes of gut microbiota composition and circulating metabolites in stroke mice. We performed 16S rRNA sequencing of the gut microbiota and serum metabolomics analysis in male mice three days post-stroke. We observed significant reduction in the alpha diversity of gut microbiota in MCAO mice (Fig. S2a), as well as notable differences in beta diversity between MCAO mice and controls (Fig. 1i). The abundances of *Lactobacillaceae* and *Clostridiales* were markedly reduced in stroke mice (Fig. 1j-k, Fig. S2b), resembling the gut dysbiosis patterns observed in PSCI patients during the acute phase. Untargeted serum metabolomics analysis identified metabolites with significant changes after stroke, including substantial reduction in tryptophan metabolites such as IPA and ILA (indole-3-lactic acid), both produced by gut bacteria^31,32^ (Fig. 1l-m). Targeted serum metabolomics analysis through LC-MS/MS further confirmed that serum levels of IPA, ILA and IAA (indole-3-acetic acid) significantly decreased three days post-stroke in mice, with more pronounced reductions compared to other tryptophan metabolites, such as KYN (kynurenic acid), 5-HTP (5-hydroxytryptophan), I3C (indole-3-carboxaldehyde), and IPY (indole-3-pyruvate) (Fig. 1n, Fig. S2c-h). Moreover, fldH, fldB, fldC and acdA, which are involved in the metabolic pathway from tryptophan to IPA^12,32^, were reduced in the gut microbiota of MCAO mice (Fig. S2i-m). These results suggest critical disruption in gut microbiota-derived tryptophan metabolites after stroke.

Strikingly, immediate intervention with IPA post-occlusion was associated with a significant reduction in infarct size, an effect not observed with either ILA or IAA (Fig. S3). In addition, the levels of IPA experienced a precipitous drop as early as the first day following stroke, with this decrease persisting up to the seventh day (Fig. 1o), indicating that acute stroke leads to a sustained reduction in serum IPA levels. To determine whether the replenishment of tryptophan-metabolizing gut bacteria reduced after stroke could recover IPA levels, we performed targeted bacterial colonization to MCAO mice with *Clostridium sporogenes* and *Lactobacillus reuteri*, two representative bacteria from the families *Clostridiaceae* and *Lactobacillaceae* respectively^32–34^. Mice colonized with *C. sporogenes* exhibited significantly elevated serum IPA levels relative to those colonized with *L. reuteri*, whereas levels of ILA and IAA remained unchanged (Fig. S4a). Furthermore, *C. sporogenes* colonization resulted in a more pronounced reduction in neuronal apoptosis compared to *L. reuteri* (Fig. S4b). These findings support that IPA may represent a key gut bacterial metabolite mediating neuroprotection after stroke.

### The rapid supplementation with IPA improves post-stroke learning and memory in MCAO mice

To investigate whether and how IPA regulates post-stroke cognitive function, RNA sequencing was performed on the ischemic hemisphere of MCAO mice with administration of IPA or vehicle at the post-stroke phase as well as ipsilateral hemisphere of sham-operated mice. Gene set enrichment analysis (GSEA) revealed a marked transcriptional shift toward cognitive function-related programs following IPA treatment. Specifically, two independent learning-and memory-associated gene sets showed strong positive enrichment in IPA-treated ischemic hemispheres (Fig. 2a-b), indicating that IPA activates the pathways relevant to cognitive processes. Moreover, gene sets involved in ligand-gated ion channel activity, AMPA receptor regulation, hippocampal CA3 synaptic activity, and voltage-gated ion channel regulation all displayed coordinated upregulation, suggesting that IPA enhances both synaptic responsivity and membrane potential dynamics disrupted after stroke. Consistent with these observations, KEGG pathway analysis of the top 10 down-regulated pathways in MCAO versus Sham mice revealed broad suppression of neurotransmission-related signaling, including neuroactive ligand-receptor interaction, calcium signaling, cAMP signaling, and glutamatergic, dopaminergic and serotonergic synapse (Fig. 2c). GO analysis further showed that MCAO induced significant reductions in chemical synaptic transmission, trans-synaptic signaling, and regulation of synaptic plasticity (Fig. 2d). Importantly, IPA administration robustly reversed these deficits (Fig. 2e-f). Several enriched modules, such as muscle system process, locomotory behavior, and response to amphetamine, further point to normalization of neuronal excitability and network activity (Fig. 2f). Collectively, these transcriptomic signatures demonstrate that IPA selectively reactivates synaptic transmission and plasticity-associated pathways that are profoundly impaired after stroke. The strong convergence across GSEA, KEGG, and GO analyses supports a mechanistic model in which IPA restores neurochemical signaling and circuit-level processes essential for cognitive recovery.

**Fig. 2.**
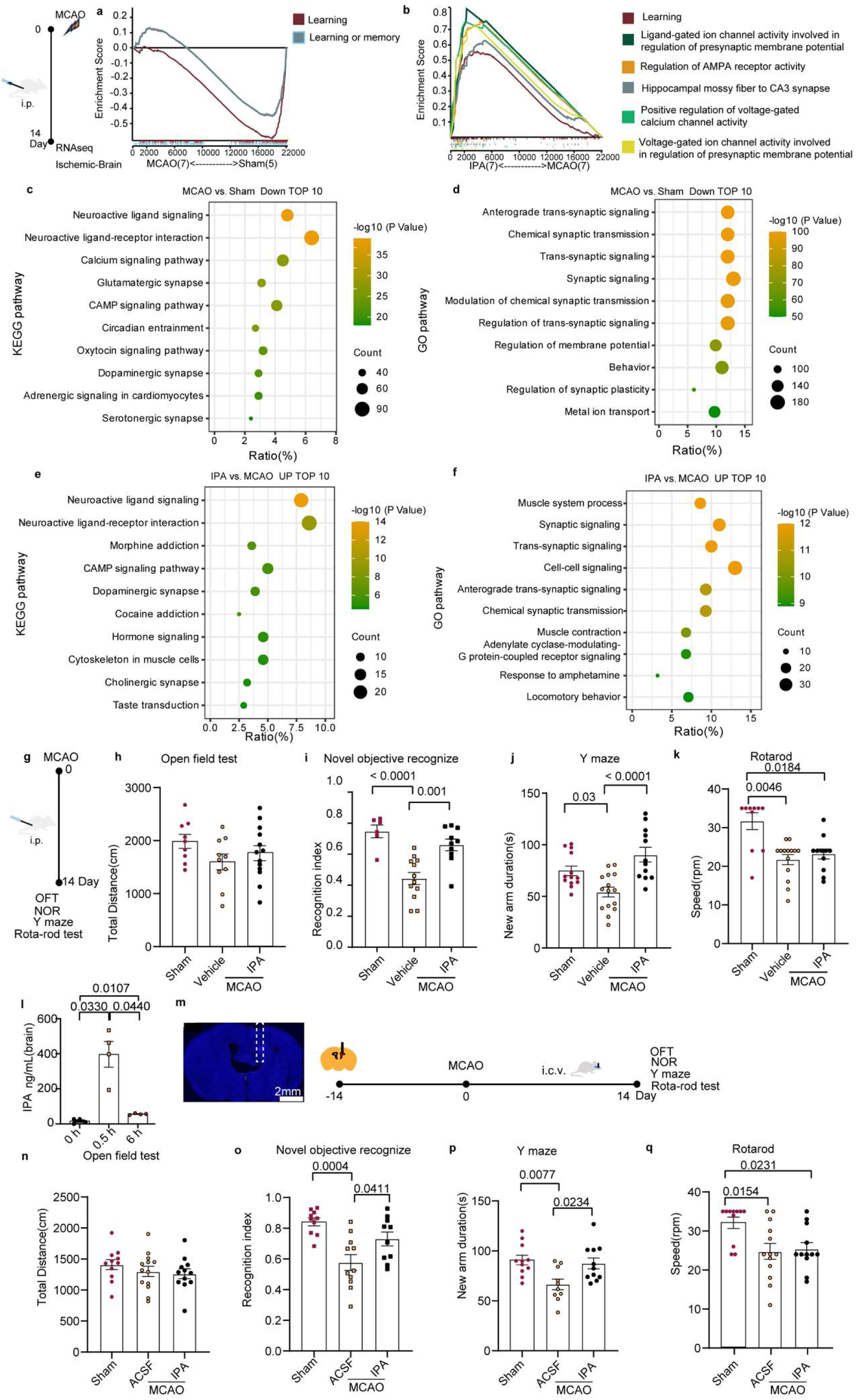
The rapid supplementation with IPA improves post-stroke learning and memory in MCAO mice. (a) Gene set enrichment analysis of MCAO group versus Sham group. (b) Gene set enrichment analysis of IPA group versus MCAO group. (c-d) Top 10 KEGG pathways (c) and GO biological processes (d) downregulated in MCAO group versus Sham group. (e-f) Top 10 KEGG pathways (c) and GO biological processes (d) upregulated in IPA group versus MCAO group. (g) The schematic diagram of MCAO mice behavioral tests following i.p. administration of IPA. (h) Travel distance in the open field test (n = 9, 10, 13). (i) Recognition index in the novel objective recognize test (n = 6, 12, 10). (j) Time spent in exploring the novel arm in the Y-maze test (n = 13, 15, 12). (k) Maximum speed to fall in the rotarod-test (n = 9, 14, 12). (l) The IPA concentration in mice brain following i.p. injection of IPA (20 mg/kg) (n = 4). (m) The schematic diagram of MCAO mice behavioral tests following i.c.v. administration of IPA. (n) Travel distance in the open field test (n = 11, 13, 12). (o) Recognition index in the novel objective recognize test (n = 10, 11, 10). (p) Time spent in exploring the novel arm in the Y-maze test (n = 11, 10, 11). (q) Maximum speed to fall in the rotarod-test (n = 11, 13, 12). ACSF, artificial cerebrospinal fluid. One-way ANOVA with Tukey’s test (h-j, n-p). Brown-Forsythe and Welch’s ANOVA test (l). Kruskal-Wallis test with Dunn’ s test (k, q). The results are expressed as means ± s.e.m.

To validate whether IPA supplementation could improve post-stroke learning and memory of mice, the MCAO mice were subjected to learning and local motor tests after 14 days of intraperitoneal (i.p.) administration of IPA or vehicle (Fig. 2g). Here, the NOR and Y-maze tasks are used to assess spatial learning and memory (Fig. S5a-b), and open field test and rotarod (Fig. S5c) are employed to test motor coordination. No differences were observed between the two groups in the open field test (Fig. 2h). In the NOR test, IPA-treated mice showed increased exploration of new objects compared to vehicle-treated mice (Fig. 2i). A similar trend was observed in the Y-maze test, where IPA-treated mice spent more time in the novel arm than vehicle-treated controls (Fig. 2j). Additionally, there was no significant difference in falling speed between the IPA and vehicle-treated groups in the rotarod test (Fig. 2k). These findings suggest that the primary benefits of IPA administration are specifically associated with the improvement of PSCI. Notably, IPA levels in brain tissue increased significantly half one hour after i.p. administration and remained elevated for up to six hours (Fig. 2l). Consistently, IPA administration via intracerebroventricular (i.c.v.) route significantly ameliorated learning and memory deficit without affecting locomotor activity or motor coordination post-stroke (Fig. 2m-q). Thus, IPA appears to exert specific cognitive benefits following stroke, independent of its effects on motor performance. However, IPA treatment through either i.p. or i.c.v. route initiated 24 hours post-MCAO did not improve NOR or Y-maze performance compared to vehicle treatment (Fig. S5d-g). These results emphasize the importance of early IPA administration in preserving cognitive function post-stroke.

### IPA ameliorates PSCI through preserving tyrosine hydroxylase-positive fibers and catecholamines levels in dorsal hippocampus

To explore the mechanisms underlying the effect of IPA on cognitive recovery, the Short Time-series Expression Miner (STEM) was employed to classify all expressed genes into eight distinct profiles (Profile 0 to 7) (Fig. 3a). Genes in Profile 2 were enriched for terms like “response to catecholamine” and “response to monoamine”, implying that IPA might be involved in the regulation of monoaminergic neurotransmitter pathways (Fig. 3b). Among these genes, tyrosine hydroxylase (Th), a key enzyme in monoamine synthesis critical for learning and memory^35,36^, was selectively upregulated by IPA post-stroke (Fig. 3c). These findings suggest that IPA may promote post-stroke cognitive recovery by regulating catecholamine. The hippocampus and mPFC (medial prefrontal cortex) are two key regions implicated in cognition and spatial memory^37,38^. In the ischemic hemisphere, the dHPC (dorsal hippocampus), unlike mPFC, exhibited significantly reduced Th fiber density compared to the contralateral side, which could be ameliorated by IPA treatment (Fig. 3d-e and Fig. S6a-b). Furthermore, the release of norepinephrine and normetanephrine from Th fibers in the ischemic dHPC was unexpectedly reduced post-stroke, which could be partially improved by IPA treatment (Fig. 3f). No notable differences in cholinergic fiber, which is responsible for acetylcholine release, learning and memory, were observed between the ipsilateral and contralateral dHPC (Fig. S6c-d). Selective ablation of Th^+^ fibers in the ipsilateral dHPC blocked IPA-mediated improvement in cognitive function (Fig. 3g). Co-administration of IPA (i.p.) during Th^+^ fiber depletion failed to protect or restore catecholaminergic projections, indicating that additional mechanisms beyond structural preservation may underlie IPA’s neuroprotective effects (Fig. S7). Together, these findings support a critical role for dorsal hippocampal Th^+^ fibers and central catecholamines in IPA-mediated improvement of cognitive function post-stroke.

**Fig. 3.**
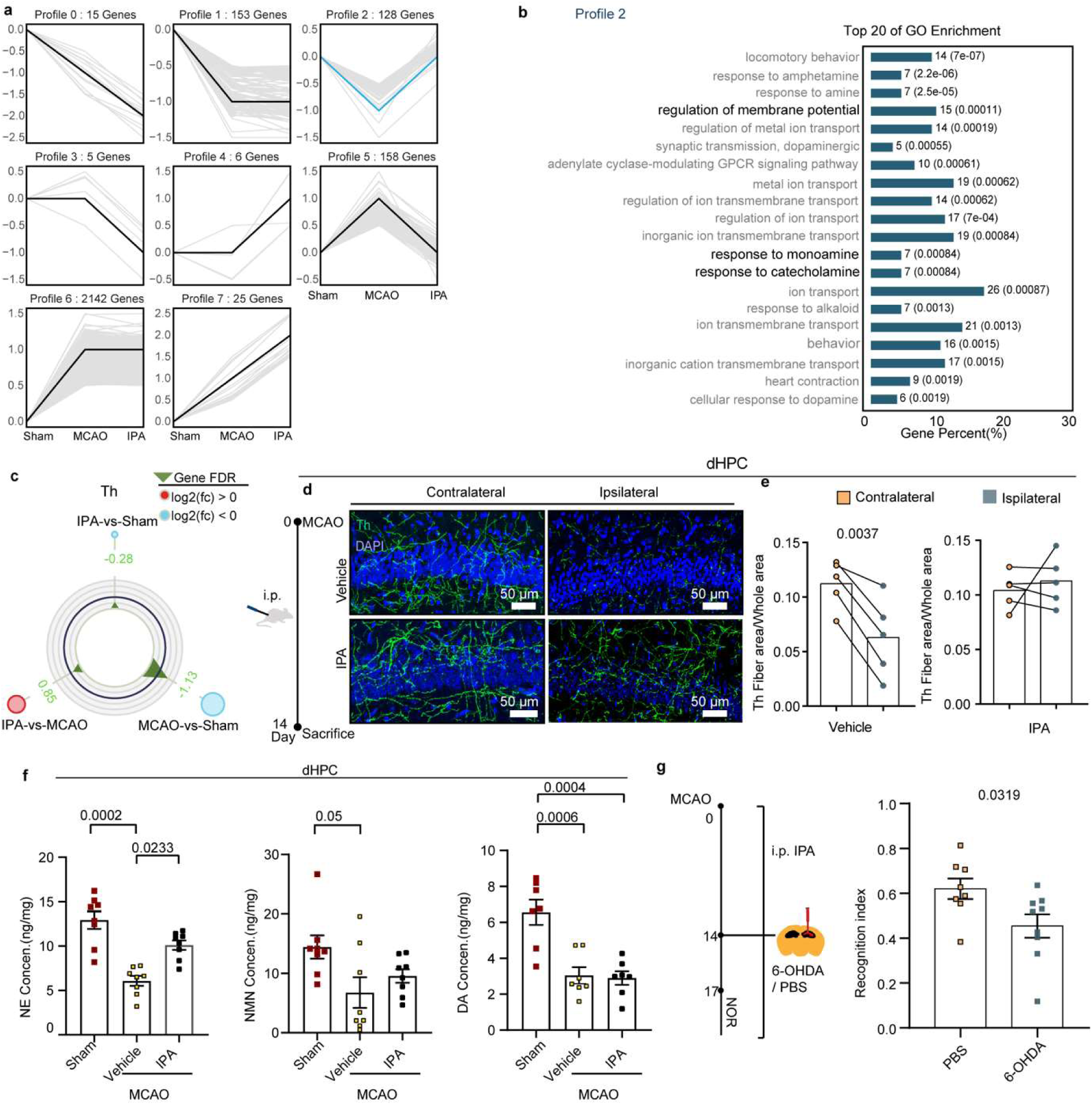
IPA ameliorates PSCI through preserving tyrosine hydroxylase-positive fibers and catecholamines levels in dorsal hippocampus. (a) Gene expression profiles categorized into clusters based on differential regulation among MCAO mice treated with IPA or vehicle and Sham (MCAO: MCAO mice treated with vehicle; IPA: MCAO mice treated with IPA). (b) Top 20 enriched pathways in Profile 2 (a). (c) The differential expressions of tyrosine hydroxylase (Th) among three groups, shown in radar plot. (d) Immunofluorescence staining of Th^+^ fibers in the contralateral and ipsilateral dorsal hippocampus in vehicle- and IPA-treated MCAO mice. (e) The quantification of Th^+^ fiber density in the contralateral and ipsilateral dorsal hippocampus (n = 5 per group). (f) The levels of catecholamines (NE, NMN and DA) in the dorsal hippocampus of Sham and vehicle- or IPA-treated MCAO mice (n = 7-8). (g) The schematic diagram showing behavior tests of MCAO mice following IPA treatment and 6-OHDA injection (left panel); New object cognitive performance of IPA-treated MCAO mice with or without 6-OHDA injection (right panel; n = 8, 9). Paired t test (e). One-way ANOVA with Tukey’s test for NE and DA; Kruskal-Wallis test with Dunn’s test for NMN (f). Mann-Whitney test (g). The results are expressed as means ± s.e.m.

Considering that IPA treatment could also led to the enrichment of pathways associated with ‘presynaptic membrane potential’, ‘voltage-gated calcium ion activity’, and ‘regulation of membrane potential’ (Fig. 2b and Fig. 3b), we next asked whether IPA directly modulates post-stroke neuronal excitability. We performed a series of experiments examining calcium signaling and local field potentials (LFP) under baseline, MCAO and reperfusion conditions. Mice were injected with AAV-hSyn-GCaMP6m in the hippocampus and fiber-optic cannulas were positioned in the injection site to allow real-time recording of calcium transients (Fig. S8a-b). Within the first hour post-IPA administration, no significant differences in calcium signal were observed between the vehicle- and IPA-treated groups, suggesting that IPA treatment did not have a marked effect on neuronal activity (Fig. S8c). The experimental protocol involved a baseline recording session, followed by i.p. administration of IPA or vehicle at 30 minutes prior to MCAO. Recordings continued through 400 seconds of baseline, followed by occlusion and subsequent reperfusion periods to assess treatment effects across ischemic stages (Fig. S8d). Under baseline conditions, calcium transients in both vehicle- and IPA-treated groups exhibited stable low-frequency oscillations (Fig. 8e, left panel). During occlusion and reperfusion, vehicle- and IPA-treated mice showed no significant differences in calcium signal (Fig. S8e, middle and right panels), indicating that IPA has no significant effect on neuronal excitability during the acute phase of ischemia-reperfusion. Next, we performed LFP recordings in the hippocampus, with electrodes positioned (Fig. S8f-g). Relative to baseline recordings, ischemic infarction and reperfusion revealed no significant differences in oscillatory power across δ, θ, α, β and low γ bands between the treatment groups (Fig. S8h-i). These findings suggest that IPA does not have significant effect on neuronal excitability or neural network activity under ischemic stress.

### IPA preserves Th^+^ fiber through inhibiting neuronal apoptosis-induced neuroinflammation in dorsal hippocampus

Next, we explored how IPA preserves Th^+^ fiber in dHPC. Post-stroke neuroinflammation, driven by damage-associated molecular patterns (DAMPs) from injured neurons, exacerbates hippocampal pathology. Here, GSEA revealed that IPA treatment negatively modulates the pathways related to immune activation and inflammation, including “the modulation of innate and adaptive immune responses”, “cytokine production involved in inflammatory response” and “microglial cell activation” (Fig. 4a). Consistently, IPA could significantly downregulate the expressions of Aif1 (a microglial marker, Iba1), Nfkbia and IL-1β in the ischemic hemisphere (Fig. 4b). Since stroke initiates immune cell infiltration and inflammatory response, leading to damage of Th^+^ fiber and catecholamine metabolism^39,40^, we investigated whether IPA could protect dorsal hippocampal Th^+^ fibers by alleviating post-stroke neuroinflammation. As expected, we found that microglial fluorescence intensities in the ischemic dHPC were significantly reduced following IPA treatment (Fig. 4c-d). Furthermore, pro-inflammatory cytokines, including TNF-α, IL-6 and IL-1β, were significantly downregulated in the infarcted hemisphere hippocampus of IPA-treated mice compared to the vehicle-treated group (Fig. 4e). Previous studies have shown that TNF-α, rather than IL-17A, IL-6 and IL-1β, directly impairs catecholaminergic nerve fibers and induces apoptosis of catecholaminergic neurons^40,41^. We therefore tested whether selective blockade of TNF-α is sufficient to mimic the protective effect of IPA on Th⁺ fibers. The result showed that continuous administration of TNF-α-neutralizing antibodies into the hippocampus post-stroke effectively preserved Th^+^ nerve fiber density (Fig. 4f-g), supporting TNF-α as a key downstream effector linking neuroinflammation to Th+ fibers loss. These data indicate that IPA protects Th⁺ fiber in dHPC at least in part by restraining TNF-α-mediated neuroinflammation.

**Fig. 4.**
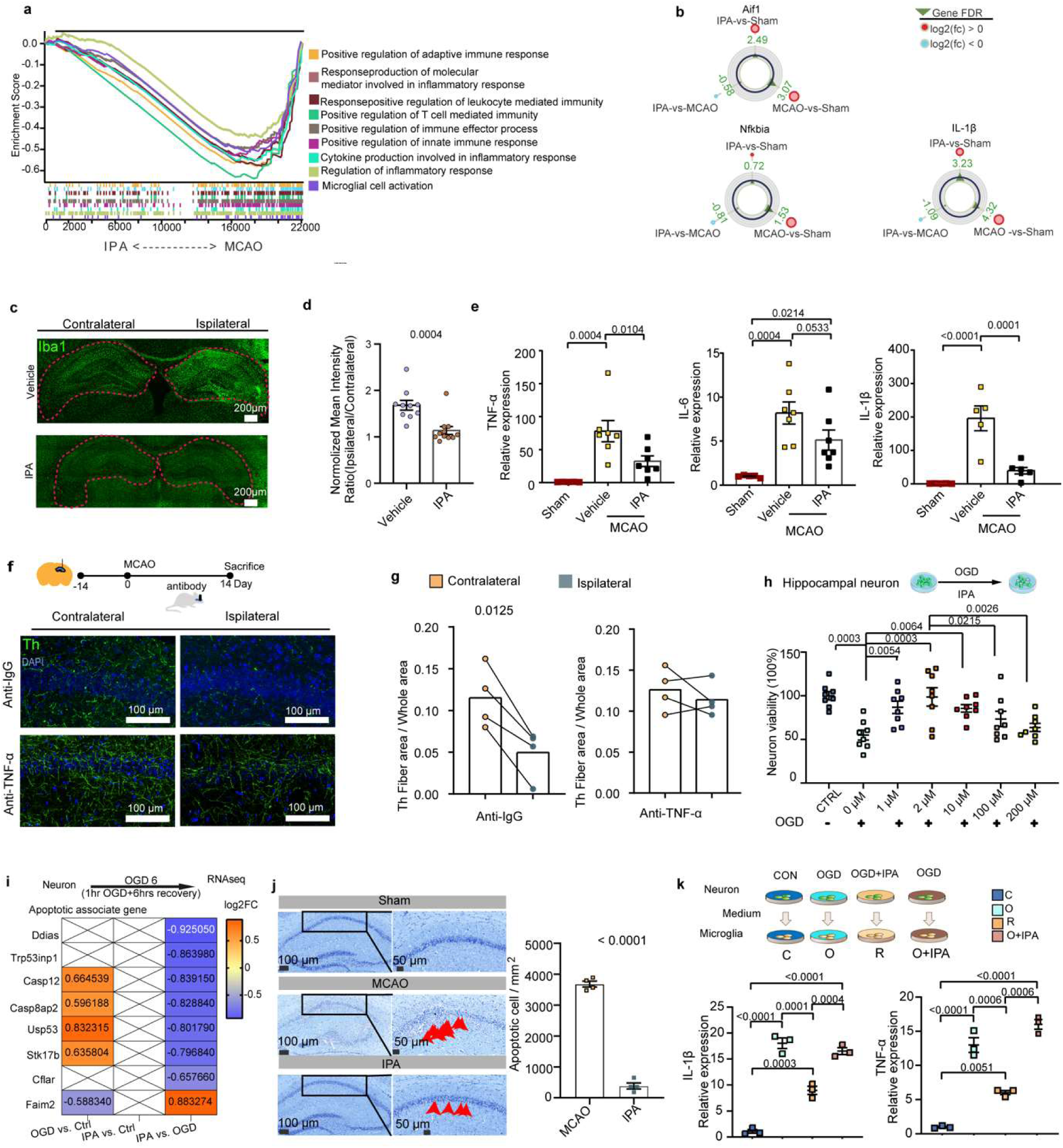
IPA preserves Th^+^ fiber through inhibiting neuronal apoptosis-induced neuroinflammation in dorsal hippocampus. (a) Gene set enrichment analysis comparing IPA-treated and vehicle-treated MCAO mice (MCAO: MCAO mice treated with vehicle; IPA: MCAO mice treated with IPA). (b) Differential expression of Aif1, Nfkbia and IL-1β among three groups, shown in radar plot. (c) Immunofluorescence staining of Iba1 in the contralateral and ipsilateral dorsal hippocampus of IPA-treated and vehicle-treated MCAO mice. (d) Relative expression of Iba1 (ipsilateral/contralateral) in the dorsal hippocampus of IPA-treated and vehicle-treated MCAO mice (n = 10, 11). (e) Relative expression of pro-inflammatory cytokines (TNF-α, IL-6 and IL-1β) in the ipsilateral dorsal hippocampus of IPA-treated and vehicle-treated MCAO mice (n = 5-7). (f) Immunofluorescence staining of Th^+^ fibers in both ipsilateral and contralateral regions of hippocampus in MCAO mice treated with Anti-IgG or Anti-TNF-α. (g) The quantification of Th^+^ fiber density in the ipsilateral and contralateral regions of hippocampus in MCAO mice treated with Anti-IgG or Anti-TNF-α, expressed as Th fiber area per whole area (n = 4 per group). (h) The anti-apoptotic effect of IPA on hippocampus neuron under oxygen-glucose deprivation (OGD) conditions (n = 8 per group). (i) The differentially expressed changes of apoptosis-related genes in OGD-exposed neurons treated with IPA or vehicle. (j) The number of apoptotic neuronal cells in the hippocampus of MCAO mice treated with IPA or vehicle (n = 4). Red arrowheads denote apoptotic neurons. (k) The expressions of IL-1β and TNF-α in BV2 microglia treated with neuron-conditioned medium under OGD conditions after differential IPA treatment (n = 3 per group). Mann-Whitney test (d). One-way ANOVA with BKY’s test (e, h). Paired t test (g). Unpaired t test (j). One-way ANOVA with Tukey’s test (k). The results are expressed as means ± s.e.m.

Then, we investigated how IPA inhibits neuroinflammation in dHPC. In primary hippocampal neurons subjected to OGD (oxygen-glucose deprivation), IPA treatment conferred dose-dependent neuroprotection, with significant anti-apoptotic effects observed at 1 μM and maximal efficacy at 2-10 μM (Fig. 4h). In order to assess the effect of IPA on gene expression in hippocampal neurons under OGD, RNA sequencing was performed on neurons exposed to OGD with or without 6-hour IPA pretreatment. Importantly, apoptosis-related genes were significantly inhibited by IPA treatment (Fig. 4i), further supporting its protective role against neuronal apoptosis. Consistently, IPA treatment markedly reduced apoptotic neurons in the hippocampus of MCAO mice (Fig. 4j), as examined by Nissl staining^42–45^. Intriguingly, co-culture of microglia with conditioned medium of OGD-exposed neurons triggered robust inflammatory responses, including elevated TNF-α and IL-1β expression, and medium from neurons under OGD plus IPA treatment (2μM) significantly attenuated microglial cytokine production, whereas simultaneous administration of IPA and medium of OGD-exposed neurons failed to replicate this suppression (Fig. 4k). These results suggest that IPA inhibits neuroinflammation in dHPC through alleviating hippocampal neuronal apoptosis. Taken together, our findings demonstrate that IPA preserves Th^+^ fiber through inhibiting neuronal apoptosis-induced neuroinflammation in dHPC.

### IPA promotes hippocampal neuronal survival via orchestrating Ywhab-ERK pathway

To elucidate the molecular mechanisms underlying IPA-mediated neuroprotection in dHPC following stroke, we employed label-free small molecule target identification techniques to identified the targets modulated by IPA. As a result, Ywhab and Gpd1 were identified as high-probability IPA-binding targets (Fig. 5a-b, Fig. S9a). Surface plasmon resonance (SPR) analyses further confirmed robust interaction between IPA and Ywhab (Fig. 5c). Functional characterization revealed that IPA directly inhibits enzymatic activity of Gpd1 *in vitro* (Fig. S9b). Spatial expression profiling via three-dimensional reconstructions and two-dimensional in situ hybridization (ISH) demonstrated prominent hippocampal enrichment of both *Ywhab* and *Gpd1* transcripts (Fig. 5d, Fig. S9c). To determine which target protein mediates the anti-apoptotic effect of IPA, shRNA-mediated knockdown of Ywhab or Gpd1 was conducted in hippocampal neurons. No significant effect on neuronal survival was observed under normal conditions following knockdown of either gene (Fig. 5e, Fig. S9d). However, knockdown of Ywhab, but not Gpd1, significantly inhibited neuronal apoptosis under OGD conditions (Fig. 5e, Fig. S9d.). Cell type-specific expression analysis confirmed the predominant localization of Ywhab in neurons instead of astrocytes or microglia (Fig. 5f). Moreover, transcriptomic analyses further revealed that OGD suppressed cAMP, calcium and MAPK signaling pathway, while IPA treatment robustly reversed this inhibition (Fig. 5g-h). Although IPA appeared to upregulate the cAMP signaling pathway, a pathway associated with long-term recovery via synaptic plasticity and angiogenesis^46^, and the calcium signaling pathway, IPA failed to restore local field potentials, which reflect synaptic activity and network-level changes associated with synaptic plasticity, and calcium dynamics in the dHPC during the acute phase of stroke (Fig. S8). These findings may not support cAMP or calcium signaling as the key pathways of IPA’s protective effect. Instead, we found that IPA significantly enhanced the phosphorylation of ERK, a key player of MAPK pathway and known to rapidly respond to ischemia-hypoxia and support neuronal survival^46^, in OGD-challenged neurons (Fig. 5i)., Moreover, Ywhab knockdown could elevate ERK phosphorylation (Fig. 5j). Interestingly, when Ywhab was knocked down, IPA failed to further enhance ERK phosphorylation (Fig. 5k), indicating that IPA promotes the activation of ERK by binding Ywhab under ischemic conditions. Collectively, these results support that IPA exerts neuroprotection by orchestrating the Ywhab-ERK signaling axis, thereby maintaining hippocampal neuronal survival during ischemic stress.

**Fig. 5.**
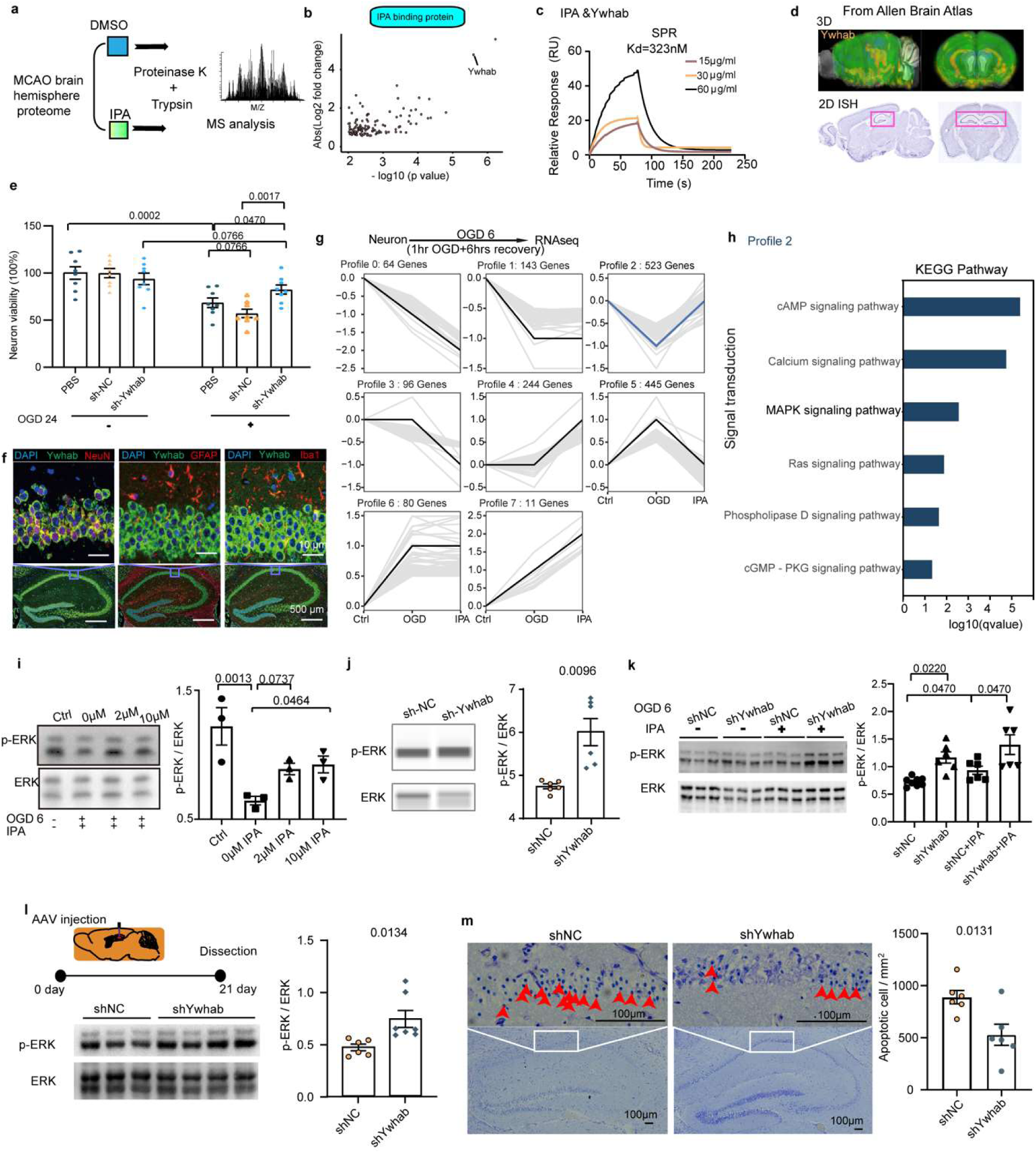
IPA promotes hippocampal neuronal survival via orchestrating Ywhab-ERK pathway. (a) Schematic diagram for label-free small molecule target protein identification technique. (b) Ywhab was identified as one of the most prominent IPA-binding target proteins in brain (n = 3 per group). (c) Surface plasmon resonance (SPR) analysis showing the binding kinetics between IPA and Ywhab, with a Kd value of 323 nM (n = 3 per group). (d) Spatial mapping of Ywhab expressions in mice brain sections from the Allen Brain Atlas. The 3D reconstruction shows strong expressions of it in hippocampus, while 2D in situ hybridization (ISH) images confirm this in corresponding brain slices. (e) The anti-apoptotic effects of Ywhab knockdown on neuron under OGD conditions (n = 8 per group). (f) The immunofluorescence co-staining of Ywhab with neuronal marker NeuN, astrocyte marker GFAP and microglia marker Iba1. (g) Gene expression profiles categorized into clusters based on differential regulation among OGD-neuron treated with IPA or vehicle and control. (h) The KEGG enrichment pathways related to signal transduction in Profile 2 (g). (i) IPA treatment promoted ERK phosphorylation in OGD-neurons (n = 3 per group). (j) The knockdown of Ywhab promoted ERK phosphorylation in neurons (n = 6 per group). (k) IPA failed to further enhanced ERK phosphorylation in neurons upon Ywhab knockdown (n = 8, 6, 6, 6). (l) The mice experimental pipeline: AAV carrying control shRNA (shNC) or Ywhab shRNA (shYwhab) was injected into the dorsal hippocampus on day 0, and the tissue was harvested on day 21. Representative immunoblot of phosphorylated ERK (p-ERK) and total ERK with quantification of the p-ERK/ERK ratio (n = 6, 7). (m) Nissl staining in the hippocampus of MCAO mice treated with shYwhab or shNC (left). Red arrowheads denote apoptotic neurons. The number of apoptotic neuronal cells were quantified (right; n = 6, 6). One-way ANOVA with BKY’s test (e, i). Unpaired t test (j, l, m). Brown-Forsythe and Welch ANOVA tests with BKY’s test (k). The results are expressed as means ± s.e.m.

To determine whether knockdown of Ywhab in hippocampal neurons could reverse the PSCI phenotype in mice, we employed an AAV2/9 viral vector system to deliver shRNA (short hairpin RNA) targeting Ywhab specifically in dorsal hippocampal neurons (Fig. 5l). In AAV-shYwhab-infected hippocampi, we firstly observed a marked increase in ERK phosphorylation (Fig. 5l), consistent with our *in vitro* findings. Behaviorally (Fig. S10a), sham-operated mice infected with AAV displayed normal cognitive performance between shNC and shYwhab groups, indicating no adverse effects of Ywhab knockdown under physiological conditions (Fig. S10b, d). Strikingly, in MCAO mice, AAV-mediated knockdown of Ywhab in dorsal hippocampal neurons significantly restored cognitive performance across two distinct learning paradigms: MACO mice with Ywhab knockdown spent more time exploring the novel object (Fig. S10c) and showed increased time in the novel arm of the Y-maze (Fig. S10e). Furthermore, in hippocampal slices from AAV-shYwhab group, we observed reduced neuronal apoptosis compared to controls (Fig. 5 m), accompanied by attenuated microglial activation (Fig. S10f-g) and preserved density of Th⁺ fibers (Fig. S10h-i). These results collectively suggest that Ywhab knockdown confers neuroprotective effects following stroke, at both molecular and behavioral levels.

## Discussion

We identify gut microbiota-derived metabolite IPA as a key regulator of cognitive recovery after stroke. Our results demonstrate that IPA activates the neuronal Ywhab-ERK signaling axis to protect hippocampal neurons, restrain microglial inflammation, and maintain the integrity of Th⁺ catecholaminergic projections, thereby supporting cognitive recovery after stroke. These findings reveal a gut-brain mechanism by which gut bacterial metabolite modulates central nervous system recovery, and establish IPA as a promising candidate for PSCI therapy.

Although hippocampal atrophy and network disconnection are well-documented in PSCI^51,52^, current therapeutic strategies rarely address these circuit-level deficits. Accelerated ipsilesional hippocampal atrophy is consistently associated with memory decline, even in the absence of direct infarction, and longitudinal imaging studies link hippocampal degeneration to poor cognitive outcomes^53^. Yet most interventions focus on cortical plasticity, with limited attention to hippocampal or neuroendocrine pathways. We demonstrate that IPA selectively preserves dorsal hippocampal Th⁺ fibers and maintains catecholamine levels, key components of the locus coeruleus-hippocampus axis essential for learning and memory^54–56^. Given the limited efficacy of current cholinesterase inhibitors, targeting this catecholaminergic circuit with IPA represents a promising therapeutic strategy. Notably, combining IPA with norepinephrine reuptake inhibitors (e.g., reboxetine or atomoxetine) may yield synergistic benefits by pairing neuronal protection with enhanced neurotransmitter availability, opening new avenues for the treatment and prevention of PSCI.

Currently, there is no validated target-driven therapy for PSCI, and available drugs remain largely symptomatic and non-specific. We define IPA-Ywhab-ERK axis as a neuron-intrinsic, mechanistically tractable pathway for intervention. IPA promotes ERK activation and neuronal survival, preventing DAMP release and secondary neuroinflammation, through binding Ywhab, a 14-3-3 family protein selectively expressed in neuron. Notably, delayed IPA treatment or post-DAMP application fails to suppress neuroinflammation, underscoring that IPA acts upstream of hippocampal inflammatory response by preserving neuronal viability. The recent identification of Ywhab homolog, Ywhag, as a candidate biomarker in Alzheimer’s disease^57,58^, further supports the IPA-Ywhab-ERK axis as potential therapeutic avenue in PSCI.

While our study identifies a dorsal hippocampal neuron-intrinsic mechanism linking gut bacterial metabolite IPA to PSCI, it captures only part of a multifactorial process. Other metabolites, brain regions, and signaling pathways likely contribute and warrant further investigation. Clinically, although serum IPA levels increased in PSCI patients, larger and longer-term cohorts are needed to validate its predictive and prognostic value. Finally, while Ywhab emerges as a promising target, its drugability remains to be confirmed through structure-based studies and *in vivo* target engagement.

In summary, the study defines a gut-brain circuit mechanism whereby gut bacteria-derived IPA promotes cognitive recovery after stroke via the neuron-intrinsic Ywhab-ERK pathway. IPA emerges as a key gut bacterial metabolite that preserves dorsal hippocampal neuronal integrity and catecholaminergic signaling to prevent PSCI. Our findings contribute to develop potential therapeutic strategies in preventing PSCI by targeting gut microbiota, IPA, and Ywhab-ERK signaling axis, and also provide promising targets for the treatment of this critical post-stroke complication. Future work should explore the broader landscape of gut microbiota-brain interactions and assess whether modulating IPA or its downstream effectors can benefit other forms of cognitive impairment beyond stroke.

## Author contributions

Conceptualization: Y.S.L., D.X.Z, H.W.Z.; Methodology: Y.S.L., H.Y.Z., F.B.X., F.W., Z.L.; Clinical investigation and analysis: J.Y., X.X.G., X.Z., M.X.L., H.Y.Z., K.Y.X., M.X.C., R.Y.R., W.K.H.; Visualization: Y.S.L., F.B.X., X.X.G.; Funding acquisition: H.W.Z., D.X.Z., F.B.X., X.X.G. Supervision: Y.S.L., D.X.Z, H.W.Z. Writing-original draft: Y.S.L., D.X.Z., H.W.Z.; Writing-review & editing: D.X.Z., H.W.Z., Y.S.L. , J.Y.

## Methods

### RESOURCE AVAILABILITY

#### Lead contact

Further information and requests for resources and reagents should be directed to and will be fulfilled by the Lead Contact, Dongxin Zhang.

#### Materials availability

All requests for resources and reagents should be directed to and will be fulfilled by the Lead Contact. All reagents including antibodies, bacteria and plasmid may be available on request after completion of a Materials Transfer Agreement.

#### Data and code availability

The raw sequence data has been deposited at the Genome Sequence Archive under accession number PRJCA034574 and are available at the following URL: https://ngdc.cncb.ac.cn/gsub/submit/bioproject/subPRO034632/overview.

For any additional information required to reanalyze the data reported in this article, please contact the primary author.

### EXPERIMENTAL MODEL AND STUDY PARTICIPANT DETAILS

#### Human study subjects

Cohort, recruited at Nanfang Hospital and Zhujiang Hospital (Southern Medical University, Guangzhou, China) (Table S1), were enrolled consecutively based on the following criteria: age ≥18 years, a confirmed diagnosis of ischemic stroke due to cerebral infarction using standard clinical criteria and imaging (CT, MRI, or MRA). Exclusion criteria included a history of major psychiatric disorders, neurodegenerative diseases, head trauma, or substance abuse. The study was approved by the Ethics Committee of Zhujiang Hospital (NO. 2021-KY-178-01). Written informed consent was obtained from all participants, and the research followed the Declaration of Helsinki principles.

#### Animals and experimental protocols

Male C57BL/6 mice were housed in standard laboratory cages, accommodating four to five mice per cage. They were maintained on a 12-hour light/dark cycle, and kept in a temperature-controlled environment ranging from 21°C to 25°C. All animals were provided with standard chow and had ad libitum access to food and water. At the time of middle cerebral artery occlusion (MCAO), the mice were 10 weeks old. All animal procedures were conducted following approval by the Animal Ethics Committee of Zhujiang Hospital, Southern Medical University. Animal experiments follow ARRIVE2.0 guidelines, including explicit reporting of study design, randomization, blinding, sample-size justification, animal characteristics, housing conditions, and ethical approvals.

#### MCAO model of mice

Following anesthesia with 1.25% tribromoethanol via intraperitoneal injection (0.02 ml/g body weight), the mouse was positioned supine on the operating table and the neck fur was shaved. A midline incision was made on the neck, and glandular tissue and fascia were dissected bluntly with forceps. Under a microscope, the common carotid artery (CCA), external carotid artery (ECA), and internal carotid artery (ICA) were exposed. The CCA was isolated and secured with a single ligature. Subsequently, the ECA and ICA were isolated distally along the CCA. The ECA and occipital artery were ligated, followed by ligation of the CCA. The proximal end of the ECA was secured with a ligature. A 45° incision was made between the two ligatures on the ECA using microsurgical scissors, and a filament was inserted into the CCA and secured with the ECA ligature. The ECA was cut proximally to grasp the ligature, aligning it with the filament in the direction of the ICA. The silicone-coated filament was carefully inserted through the ECA into the ICA until it reached the origin of the middle cerebral artery. The mouse was placed in an incubator for 60 minutes, after which the filament was gently withdrawn. The surgical site was disinfected, and the incision was sutured to complete the procedure. Exclusion criteria included: (1) mouse mortality during the procedure, or (2) absence of brain ischemia as confirmed by histological analysis. At the study’s conclusion, mice were anesthetized and euthanized. Blood samples were collected and centrifuged at 3000 rpm and 4 °C for 10 min to obtain serum samples. The thoracic cavity was opened to expose the heart for perfusion and fixation with saline. The entire brain and ileum were dissected; some sections were snap-frozen in liquid nitrogen and stored at -80 °C, while others were fixed in paraformaldehyde for subsequent histological analysis.

#### Bacterial culture and *in vivo* colonization in mice

*Lactobacillus reuteri* (from our lab) and *Clostridium sporogenes* (strain 15579) were anaerobically cultured on blood agar plates. After incubation for 48 h, bacterial colonies were harvested for *in vivo* colonization experiments in mice.

Mice received oral gavage of four-antibiotic cocktail (vancomycin 100 mg/kg/day; neomycin, metronidazole, and ampicillin each at 200 mg/kg/day), dissolved in sterile water at 200 µL/day, for 7 consecutive days. This antibiotic pretreatment was designed to deplete the resident gut microbiota prior to colonization study. Afterward, mice were gavaged with *L. reuteri* or *C. sporogenes* (1 × 10⁹ CFU) according to the schematic diagram in Fig. S4a.

#### Intraperitoneal injection of drugs in mice

IPA, IAA, or ILA were administered daily via intraperitoneal (i.p.) injection starting the day after MCAO at a dose of 400 μg per 20 g of body weight^6,7,31^

#### Microinjection of drugs into the lateral ventricle of mice

A 5 μL of IPA (10 μM, dissolved in Artificial Cerebrospinal Fluid) or ACSF were injected into the lateral ventricles of free-moving mice at 0.5 μL/min, respectively after MCAO using microinjection Pump (RWD)^59^.

Each animal received 1 μL solution of 2 μg anti-IgG (BE0088, Bio X Cell ) or 2μg of purified neutralization antibodies against TNF-α (BE0058, Bio X Cell ) injected into the hippocampi^60^. Each animal received 1 μL solution of 1 μg 6-OHDA (Sigma, H4381-100mg) or vehicle injected into the hippocampi.

#### Primary neuron culture and oxygen-glucose deprivation^61^

Primary hippocampal neurons were obtained from neonatal C57BL/6 mice. Briefly, hippocampal tissues were isolated and rinsed three times in calcium- and magnesium-free Hank’s balanced salt solution (HBSS). The tissues were then enzymatically digested with papain (0.5 mg/mL), prepared in PBS supplemented with DNase I (10 μg/mL), for 15 min at 37°C. Enzymatic activity was terminated by washing the tissue with HBSS containing 10% fetal bovine serum. Subsequently, the samples were transferred into Neurobasal medium (Gibco) supplemented with B27 (1:50, Gibco), gently triturated to obtain a single-cell suspension, and seeded into poly-D-lysine-coated 96-well plates (0.1 mg/mL; Sigma-Aldrich) at a density of 3 × 10⁴ cells per well.

Cells were maintained at 37°C in a humidified atmosphere containing 5% CO₂ for 7 days prior to experimentation. For oxygen–glucose deprivation (OGD), neuronal cultures were incubated in glucose-free DMEM (Biosharp) and placed in an anaerobic chamber (Thermo Forma 1029, Thermo Fisher Scientific) under hypoxic conditions at 37°C for the indicated durations. Control cells were maintained in DMEM under normoxic conditions for 1 h.

Following OGD exposure, the culture medium was replaced with conditioned medium, and cells were returned to normoxic conditions for a 24-h recovery period. Indole-3-propionic acid (IPA; 0–200 μM) was administered during both the OGD treatment and subsequent recovery phase.

Cell viability was evaluated using the CCK-8 assay (GLPBIO) according to the manufacturer’s instructions. Briefly, cells were washed twice with PBS and incubated with fresh medium containing 10% CCK-8 reagent at 37°C for 1 h, after which absorbance at 450 nm was measured using a microplate reader.

For RNA sequencing, primary neurons were treated with IPA (2 μM) or vehicle (DMSO) for 6 h, followed by OGD or normoxic conditions. Total RNA was immediately extracted using the RNeasy Kit (Qiagen) in accordance with the manufacturer’s protocol to ensure RNA integrity for downstream transcriptomic analysis.

For gene knockdown experiments, neurons were cultured for 5 days in vitro prior to lentiviral transduction with constructs expressing shRNA targeting scramble control, Gpd1, or Ywhab (titer: 7 × 10⁸ TU/mL). Five days after infection, cells were subjected to 1 h of OGD followed by 24 h of recovery, and cell viability was subsequently assessed using the CCK-8 assay.

#### BV2 microglia cell culture and treatment^61^

BV2 microglial cells were maintained in high-glucose DMEM supplemented with 10% fetal bovine serum under standard culture conditions (37°C, 5% CO₂, humidified atmosphere). Cells were used for subsequent experiments upon reaching approximately 80% confluence, at which point the culture medium was replaced with neuron-conditioned medium to enable cell stimulation.

Total RNA was isolated using TRIzol reagent (Gibco Invitrogen) following the manufacturer’s protocol. RNA integrity was assessed using an Experion automated electrophoresis system (Bio-Rad), and concentrations were quantified with a NanoDrop 2000 spectrophotometer (Thermo Fisher Scientific). Purified RNA samples were stored at −80°C prior to downstream analyses.

For conditioned medium preparation, primary neurons were subjected to oxygen–glucose deprivation (OGD) in the presence or absence of indole-3-propionic acid (IPA, 2 μM) for 24 h, with untreated neurons maintained under normoxic conditions serving as controls. Following treatment, culture supernatants were collected and centrifuged at 40,000 × g for 1 h to eliminate cellular debris. The clarified supernatants were subsequently harvested for further analyses.

### METHOD DETAILS

#### TTC staining technique for delineating infarct zones^62^

Infarct size was assessed 3 days after middle cerebral artery occlusion (MCAO). Mice were deeply anesthetized and transcardially perfused with 10 mL of ice-cold phosphate-buffered saline (PBS). Brains were rapidly harvested and snap-frozen on dry ice to preserve tissue integrity. Coronal brain sections (1 mm thickness) were subsequently prepared for infarct visualization.

For histological assessment, sections were incubated in 2,3,5-triphenyltetrazolium chloride (TTC; Sigma-Aldrich) solution at room temperature for 15 min, followed by fixation in 4% paraformaldehyde for 30 min. Stained sections were imaged, and infarct areas were quantified using ImageJ software.

Infarct volume was calculated by integrating the infarct area across serial sections. To minimize the influence of cerebral edema, an established correction method was applied. The infarct ratio was expressed as the percentage of infarcted volume relative to the total brain volume for each animal.

### 16s rRNA sequencing and analysis

Bacterial genomic DNA was isolated utilizing the MinkaGene Stool DNA Kit in accordance with the guidelines provided by the manufacturer. Amplification of the V4 variable region of the 16S rRNA gene was achieved using barcoded primers V4F (GTGYCAGCMGCCGCGGTAA) and V4R (GGACTACNVGGGTWTCTAAT).

Subsequently, all PCR products were pooled and sequenced on an Illumina iSeq 100 system following the manufacturer’s instructions. In the processing of microbial 16S rRNA gene sequencing data, rarefaction is utilized to equalize sequencing depth across samples, effectively reducing the influence of variable sequencing efforts. In this study, sequencing data were normalized by rarefying each sample to a consistent depth of 7000 reads. This normalization strategy was selected to facilitate comparability among samples by standardizing the count of sequencing reads analyzed per sample, thereby reducing bias due to variations in sequencing output. The rarefaction was executed using the QIIME2 software suite, designed to enhance the reliability and reproducibility of microbial community analyses. Linear discriminant analysis effect size (LEfSe) was employed to delineate discriminative features among groups. This algorithm excels in high-dimensional biomarker discovery, pinpointing genomic attributes that distinguish between two or more biological states. LEfSe emphasizes statistical significance, biological consistency, and relevance of effects, allowing for the identification of features that show significant abundance differences between conditions aligned with biologically meaningful categories. For each distinct feature identified by LEfSe, a linear discriminant analysis score was calculated to quantify the extent of differentiation between groups, highlighting the most distinctive attributes. All analyses were conducted on the CALM-based Microbiome Analysis Platform (<u>CMAP</u>).

#### Bacterial quantification by quantitative PCR (qPCR) ^63^

Experimental procedures were performed based on established protocols with appropriate modifications. Fecal microbial DNA was obtained using a silica column–based extraction system (MolPure, Yeasen Biotechnology), with procedures carried out under the supplier’s guidelines. The quantity and purity of the extracted DNA were evaluated by ultraviolet spectrophotometry.

To assess the abundance of functional genes involved in indole-3-propionic acid (IPA) biosynthesis, quantitative PCR was performed using a SYBR Green fluorescence-based system (Accurate Biology). Gene-specific primers targeting IPA-associated microbial pathways are detailed in the Materials section.

#### qPCR for mRNA expressions

Quantitative analysis of mRNA expression levels was conducted for genes involved in inflammation and cellular responses, including Gapdh, IL-1β, TNF-α and IL-6 via quantitative real-time PCR (qRT-PCR). Colon tissue samples, stored in TRIzol reagent (Invitrogen), were homogenized at 70 Hz for 2 minutes using a tissue homogenizer (JXFSTPRP-32) equipped with sterile steel balls. RNA was extracted following the manufacturer’s protocol from Thermo Fisher Scientific, and cDNA synthesis was conducted using reverse transcription reagents from Takara. The SYBR Premix Ex Taq™ II (Takara) was utilized for real-time PCR, and the ViiA 7 system (Takara) captured the data. Relative transcription levels were determined using the comparative Ct method, normalizing each target gene’s transcription against Gapdh mRNA levels.

#### RNA sequencing

Transcriptomic profiling was performed on ischemic brain tissues collected from mice subjected to 14 days of indole-3-propionic acid (IPA; 20 mg/kg/day) or vehicle treatment following middle cerebral artery occlusion (MCAO). Total RNA was isolated using a silica membrane–based purification system (RNeasy, Qiagen), and RNA integrity was preserved for downstream library construction.

Sequencing libraries were generated using a strand-specific RNA library preparation workflow (NEBNext Ultra, New England Biolabs). Briefly, polyadenylated RNA was fragmented and reverse transcribed to generate first- and second-strand cDNA, followed by end repair, adaptor ligation, and amplification. Libraries were purified and subjected to high-throughput sequencing on an Illumina NovaSeq 6000 platform.

To characterize transcriptional alterations associated with ischemic injury and IPA intervention, differential expression analysis was conducted using predefined statistical thresholds (|log₂ fold change| ≥ 0.5, adjusted p < 0.05). Genes meeting these criteria were considered differentially expressed and subjected to downstream functional annotation.

Functional enrichment analyses were performed to interpret biological significance. Gene Ontology (GO) terms and Kyoto Encyclopedia of Genes and Genomes (KEGG) pathways were analyzed using the Metascape platform. In parallel, gene set enrichment analysis (GSEA) was carried out with curated gene sets from the MSigDB database to evaluate coordinated transcriptional changes at the pathway level. Gene ranking was based on signal-to-noise metrics, and enrichment significance was assessed using standard parameters.

To further resolve dynamic expression patterns, clustering analysis was conducted using the Short Time-series Expression Miner (STEM, version 1.3.11), enabling the identification of gene modules with similar expression trajectories. Genes within each cluster were subsequently subjected to functional enrichment analysis to define their associated biological processes.

All bioinformatic analyses were integrated and performed using the OmicShare platform.

#### Metabolite protein binding profiling

Proteomic analysis was performed on ischemic brain tissues to identify protein alterations associated with indole-3-propionic acid (IPA) treatment. Brain tissues were lysed under non-denaturing conditions, and clarified lysates were obtained by high-speed centrifugation. Protein concentrations were determined prior to downstream processing. Lysates were then incubated with IPA (100 μM) or vehicle control to assess ligand-associated proteome changes. Protein samples were subjected to limited proteolysis followed by enzymatic digestion. Briefly, lysates were treated with Proteinase K under controlled conditions and subsequently denatured. Proteins were reduced and alkylated, followed by trypsin digestion at a defined enzyme-to-substrate ratio to generate peptides suitable for mass spectrometry analysis. The resulting peptides were acidified, cleared of detergents, and desalted prior to chromatographic separation. Peptide mixtures were separated using a nano-flow ultra-high-performance liquid chromatography system (EASY-nLC 1000) with a multi-step acetonitrile gradient and introduced into a high-resolution mass spectrometer (Orbitrap Q Exactive) via a nano-electrospray ionization source. Data were acquired in a data-dependent acquisition (DDA) mode, in which the most abundant precursor ions from each full scan were selected for fragmentation and subsequent tandem MS analysis. Raw MS data were processed using MaxQuant (version 1.6) against a curated protein database with decoy sequences to control false discovery rates. Standard search parameters were applied, including trypsin specificity, defined mass tolerances, and common fixed and variable modifications. Protein identification was filtered at a 1% false discovery rate threshold.

#### Surface plasmon resonance (SPR) analysis of IPA-Ywhab interaction

Indole propionic acid is immobilized onto a LifeDiscm photo-crosslinking chip (Liangzhun medical technology Co., LTD, Shanghai, China). The Ywhab standard is diluted in HBS-ET buffer (pH 7.4) to final concentrations of 0 μg/mL, 15 μg/mL, 30 μg/mL, and 60 μg/mL. Using the WeSPR™ one molecular interaction analyzer (Liangzhun medical technology Co., LTD), 100 μL of 1% casein is injected first, followed by the steps of baseline establishment, analyte binding, dissociation, and regeneration with the following solutions in sequence: HBS-ET buffer (pH 7.4), Ywhab solution, HBS-ET buffer (pH 7.4), and hydrochloric acid-glycine (pH 2.0). Different concentrations of Ywhab were flowed through the LifeDiscm chip with a binding time of 150 s and a dissociation time of 300 s, followed by analysis of the Kd value. **IPA effects on GPD1 enzyme activity**

The GPD1 enzyme activity assay utilized a 100 μL reaction mixture containing 20 mM TEA buffer, 0.1 mg/mL BSA, 200 μM NADH, 8 mM DHAP, 1 μM GPD1, and either the compound or DMSO at an ionic strength of 0.12 (NaCl). The initial velocity of DHAP reduction was tracked by measuring the decrease in NADH concentration. Absorbance at 340 nm was recorded using a M2e microplate reader (Molecular Devices).

#### Western blot

Lysates were prepared from primary hippocampal neurons exposed to 6 hours of oxygen-glucose deprivation (OGD) under treatment with IPA or vehicle, as well as under normal conditions. Neurons were lysed in radioimmunoprecipitation assay (RIPA) buffer at 4°C for 30 minutes. The lysates were then centrifuged at 12,000× g for 30 minutes to remove cellular debris. Protein concentrations were determined using a BCA Protein Assay Kit. A total of 30 µg of protein from the neuronal samples was loaded onto 10% pre-cast gels. After electrophoresis, proteins were transferred to 0.2 µm PVDF membranes. To prevent nonspecific antibody binding, the membranes were blocked with 5% bovine serum albumin (BSA). The primary antibody, rabbit IgG anti-pERK (1:500, 12638, CST) was incubated overnight at 4°C with gentle shaking (90 rpm). Following the incubation, the membranes were washed four times with TBST (Tris-buffered saline with 0.05% Tween) for 10 minutes each. The secondary antibody, HRP-conjugated goat anti-rabbit IgG, was applied for 2 hours at room temperature while the membranes were rocked. After further washing, the protein signals were visualized using the Bio-Rad ChemiDoc Imaging System.

To investigate ERK phosphorylation, membranes were first stripped and then re-probed with an antibody specific to ERK protein. The membranes underwent incubation in stripping buffer at room temperature for 10 minutes with gentle agitation. Afterward, the membranes were rinsed under running distilled water (mH2O) for 2 minutes, followed by washing with TBST, as described previously. Post-stripping, the membranes were blocked and subsequently immunoblotted using rabbit anti-ERK antibody (1:200, 11257-1-AP, Proteintech) and imaged.

#### JESS Simple Western™

Lysates from primary hippocampal neurons infected with lentivirus-shYwhab for 3 days were collected for analysis. Protein concentrations were adjusted to 3 μg per well. For immunodetection, primary antibodies-rabbit polyclonal anti-pERK (1:500, 12638, CST) and rabbit polyclonal anti-ERK (1:200, 11257-1-AP, Proteintech)-were diluted in Milk-Free Ab diluent (Bio-Techne), and 10 µL of each antibody solution was added to the wells. All steps, including antibody dilutions, washing, and ECL reagent application, were performed following the manufacturer’s protocol. To enable sequential detection of both primary rabbit antibodies, the RePlex™ reagent kit was employed for dual immunodetection.

#### Immunoblot and Jess quantification

Densitometric analysis of conventional Western blot was performed using Image Lab software (Bio-Rad) with automatic lane background subtraction. Capillary-based Simple Western assays run on the Jess platform (ProteinSimple) were quantified using Compass for Simple Western software according to the manufacturer’s instructions.

#### Open field test

The open field test (OFT) was performed as previously described with slight modifications. Each mouse was placed in the center of a square box (40 × 40 × 40 cm) and allowed to explore freely for 5 minutes while the travel distance was measured. The arena was cleaned with 75% ethanol between trials.

#### Rotarod test

The mice should acclimate to the testing environment for at least 30 minutes and be handled gently to minimize stress. Training sessions familiarize the rodents with the apparatus by placing them on the stationary rod and gradually increasing the rod’s rotation speed over 1-2 days. During testing, place the mouse on the rotating rod, when conducting experiments using a small animal rotarod fatigue apparatus, the total experiment duration is set to 5 minutes. The experiment begins with an initial speed of 5 rpm/min for 20 seconds. The speed then accelerates to a primary speed of 12 rpm/min, maintained for 60 seconds. Next, the rod speed increases to 24 rpm/min, sustained for another 60 seconds. Finally, the speed accelerates to 35 rpm/min and is maintained for 40 seconds. Ultimately, the speed at which the mice fall is calculated. Afterward, the average speed when the mouse dropped down three times during training was recorded for statistical analysis. To avoid discrimination of the objects based on odor, the apparatus was thoroughly cleaned with 75% ethanol before and after each trial.

#### NOR test

The NOR test comprises three distinct phases: On day 1, a 5-minute habituation phase to familiarize the mouse with the testing arena without objects for three times; 24-hour retention interval to allow for memory consolidation of the testing arena. On day 2, a 10-minute training phase where the mouse interacts with the two same objects; On day 3, a 5-minute testing phase where one familiar object is replaced with a novel object to measure the mouse’s preference for exploring the novel versus familiar object. Recognition memory was assessed using the recognition index, calculated as follows: (time spent on the new object/total time spent exploring both objects) × 100%. To maintain cleanliness, both the arena and the objects was cleaned with a 75% ethanol solution between trials.

#### Y-maze test

Spatial working memory was evaluated using a three-arm maze constructed from opaque white plastic. Each arm (33 cm × 10 cm × 11 cm) was arranged at 120° relative to the others. The task consisted of a training session followed by a delayed test to assess novelty preference.

During the training session, one arm was temporarily closed, allowing animals to explore the two accessible arms freely for 10 min. After a 4-h inter-trial interval, the blocked arm was reopened, and animals were reintroduced into the maze from a randomized starting arm. Exploratory behavior was recorded for 3 min to determine arm preference.

To minimize olfactory cues, the maze was thoroughly cleaned with 75% ethanol between trials.

#### Immunostaining and imaging

Immunofluorescence staining was performed to characterize cellular and molecular changes in the brain following experimental treatments. Mice were deeply anesthetized and transcardially perfused with saline followed by 4% paraformaldehyde (PFA). Brains were harvested, post- fixed overnight at 4°C, and cryoprotected in graded sucrose solutions prior to sectioning. Coronal sections (40 μm) were prepared using a cryostat (Leica CM1900). After permeabilization and blocking in a bovine serum albumin–based buffer containing detergent, free-floating sections were incubated with primary antibodies against neuronal, glial, and target proteins (including Iba1, NeuN, GFAP, TH, ACh, and Ywhab) at 4°C overnight.

Following primary incubation, sections were treated with species-appropriate fluorescent secondary antibodies and subsequently mounted using an antifade medium containing DAPI for nuclear counterstaining. Fluorescence images were acquired using a laser-scanning confocal microscope (Nikon AxR) under identical acquisition settings across groups.

#### Nissl Staining

Nissl staining was performed according to the manufacturer’s instructions. The mice brains were sliced (10 μm) by a rapid sectioning cryostat (CM1900, LEICA, Germany). Begin by fixing the samples in 4% paraformaldehyde for over 10 minutes, followed by a 2-minute wash in distilled water and another 2-minute wash in fresh distilled water. Subsequently, stain the samples with Nissl staining solution for 3 minutes. After staining, perform two brief rinses in distilled water, each lasting a few seconds. Then, dehydrate the samples in 95% ethanol for about 5 seconds; if direct observation is required, rinse twice in 70% ethanol. For dehydration, transparency, and mounting, continue with 95% ethanol for 2 minutes, followed by 5 minutes in xylene, and repeat with fresh xylene for an additional 5 minutes. Finally, mount the samples using neutral gum or another mounting medium. Upon microscopic examination, cells should exhibit a mottled blue-purple stain.

#### Untargeted ultra-performance liquid chromatography-mass spectrometry

Metabolic alterations associated with experimental intervention were assessed using untargeted serum metabolomics. Serum samples were processed under cold conditions to preserve metabolite stability, followed by protein removal through organic solvent precipitation. The resulting extracts were concentrated and reconstituted for instrumental analysis. A pooled quality control sample was prepared from all experimental groups and injected periodically to ensure analytical reproducibility across the run sequence.

Chromatographic separation and metabolite detection were carried out using a high-resolution LC–MS platform. Metabolites were resolved on a reversed-phase column under gradient elution conditions and analyzed in both ionization modes to maximize metabolome coverage. Data-dependent acquisition was applied to obtain both precursor and fragment ion information for downstream annotation.

Raw spectral data were processed by matching accurate mass, retention behavior, and MS/MS fragmentation patterns against curated reference databases. Statistical evaluation of metabolic differences between groups was performed using multivariate modeling in R. Group separation and treatment-associated metabolic shifts were assessed using OPLS-DA, with model robustness validated through permutation testing. Discriminative metabolites were selected based on combined thresholds of multivariate importance and statistical significance.

#### LC-MS/MS

Targeted quantification of serum tryptophan-related metabolites was performed using LC–MS/MS analysis. A panel of standards including IPA, IAA, IPY, ILA, I3C, 5-HTP, and KYN was used for calibration and identification. Calibration curves for IPA were constructed over a defined concentration range to ensure quantitative reliability.

Chromatographic separation was achieved using a UPLC system coupled to tandem mass spectrometry. Metabolites were resolved on a reversed-phase C18 column under a rapid gradient elution using aqueous formic acid and acetonitrile as mobile phases. Samples were maintained at low temperature during automated injection to minimize degradation.

Mass spectrometric detection was performed in multiple reaction monitoring (MRM/SRM) mode using a heated electrospray ionization source under optimized instrumental conditions. Standard MS parameters, including gas flows, ion transfer temperature, and collision settings, were applied to ensure analytical stability and sensitivity.

Catecholamine levels in the dorsal hippocampus were quantified using a derivatization-assisted LC–MS/MS approach. Frozen tissue samples were homogenized, and standards and quality control samples were equilibrated to room temperature prior to processing.

Analytes were chemically derivatized using dansyl chloride following protein precipitation with acetonitrile. After sequential pH adjustment and liquid–liquid extraction, metabolites were enriched, concentrated under nitrogen, and reconstituted in a formic acid–acetonitrile aqueous solution prior to analysis.

Separation and detection were performed using LC–MS/MS, and derivatized catecholamines were quantified based on internal standard calibration. Instrumental analysis was conducted under controlled temperature and centrifugation conditions to ensure reproducibility across samples.

#### Stereotaxic surgery, virus injection, and optic fiber and cannula implantation

In vivo calcium imaging of hippocampal neuronal activity was performed using a fiber photometry system. Male mice (6 weeks old) were anesthetized and secured in a stereotaxic apparatus for viral delivery. AAV vectors encoding GCaMP6m were injected into the right dorsal hippocampus using stereotaxic coordinates, allowing region-specific expression of the calcium indicator. Following viral infusion, the injection needle was left in place briefly to facilitate diffusion.

After viral expression, optical fibers were implanted above the dorsal hippocampus for fluorescence signal collection. In a subset of experiments, guide cannulae were additionally implanted for intracerebral drug administration. All implants were secured to the skull using anchoring screws and dental cement to ensure stability for longitudinal recordings.

Following a recovery and expression period, calcium signals were recorded in freely behaving mice using a dual-wavelength fiber photometry system (470 nm for GCaMP6m-dependent activity and 405 nm for motion and bleaching correction). Fluorescence signals were recorded during behavioral testing or MCAO procedures, and animals received either indole-3-propionic acid (IPA) or vehicle treatment.

For region-specific gene manipulation, AAV-shYwhab or control virus was stereotaxically delivered into the dorsal hippocampus, and subsequent experiments were performed after sufficient expression time to ensure effective knockdown.

Signal processing and ΔF/F calculation were performed using custom MATLAB scripts.

#### EEG electrode installation and recording

Isoflurane anesthesia was administered prior to fixing mice in a stereotaxic apparatus for electrode implantation. The recording electrode was anchored to the left parietal bone (coordinates: AP -1.8 mm; ML 1.5 mm; DV 1.25 mm), the reference electrode to the right frontal bone, and the ground wire to the left occipital bone. All surgical procedures were performed under strict sterile conditions, and mice were given three days to recover before further experimental procedures.

Electroencephalographic (EEG) recordings were conducted using a Solar system (model 1848, Solar, China) at a sampling rate of 1000 Hz, with a band-pass filter ranging from 0.01 to 70 Hz. To assess the power spectral density (PSD) of the EEG signals, MATLAB’s Pwelch function was employed. Analysis included frequency bands of delta (0.5-4 Hz), theta (4-8 Hz), alpha (8-12 Hz), beta (12-30 Hz), and gamma (30-70 Hz).

#### Statistical analysis

The results were presented as mean ± standard error of mean (s.e.m.) of multiple independent experiments. Statistical analyses were performed using GraphPad Prism version 8.0. Data were tested for normality and homogeneity of variance. For parametric data or nonparametric data, comparisons between groups were performed by employing different statistical test followed by appropriate post-hoc test as indicated in the figure legends. A value of *p*<0.05 was considered statistically significant.

## Supplemental information

Supplemental Tables (Table S1-S3).

Supplemental Figures (Figure S1-S10).

